# Organ-Specific Fuel Rewiring in Acute and Chronic Hypoxia Redistributes Glucose and Fatty Acid Metabolism

**DOI:** 10.1101/2022.08.25.505289

**Authors:** Ayush D. Midha, Yuyin Zhou, Bruno B. Queliconi, Alec M. Barrios, Cyril O. Y. Fong, Joseph E. Blecha, Henry VanBrocklin, Youngho Seo, Isha H. Jain

**Affiliations:** Gladstone Institutes, San Francisco, CA, 94158, USA; Department of Biochemistry and Biophysics, University of California, San Francisco, CA 94158, USA; Medical Scientist Training Program, University of California, San Francisco, CA 94143, USA; Tetrad Graduate Program, University of California, San Francisco, San Francisco, CA 94158, USA; Department of Radiology and Biomedical Imaging, University of California, San Francisco, CA 94107, USA

## Abstract

Oxygen deprivation can be detrimental. However, chronic hypoxia is associated with decreased incidence of metabolic syndrome and cardiovascular disease in high-altitude populations. Previously, hypoxic fuel rewiring has primarily been studied in immortalized cells. Here, we describe how systemic hypoxia rewires fuel metabolism to optimize whole-body adaptation. Acclimatization to hypoxia coincided with dramatically lower blood glucose and adiposity. Using *in vivo* fuel uptake and flux measurements, we found that organs partitioned fuels differently during hypoxia adaption. Acutely, most organs increased glucose uptake and suppressed aerobic glucose oxidation, consistent with previous *in vitro* investigations. In contrast, brown adipose tissue and skeletal muscle became “glucose savers,” suppressing glucose uptake by 3-5-fold. Interestingly, chronic hypoxia produced distinct patterns: the heart relied increasingly on glucose oxidation, and unexpectedly, the brain, kidney, and liver increased fatty acid uptake and oxidation. Hypoxia-induced metabolic plasticity carries therapeutic implications for chronic metabolic diseases and acute hypoxic injuries.

## INTRODUCTION

Oxygen deprivation contributes to several leading causes of mortality in developed nations— myocardial infarction (MI), stroke, and respiratory failure. While oxygen deprivation can have devastating health consequences, chronic hypoxia can stimulate protective adaptations. For example, a gradual ascent to altitude reduces the incidence of altitude sickness, high-altitude pulmonary edema (HAPE), and high-altitude cerebral edema (HACE) (Imray et al., 2011). At the organ level, exposure to transient, sublethal ischemia increases resilience to future ischemic events, known as ischemic preconditioning (Carr et al., 1997; Lankford et al., 2006; Liu and Downey, 1992; Murry et al., 1986; Schott et al., 1990). Moreover, ischemic injury to non-cardiac tissue confers resilience to MI, known as remote ischemic preconditioning (Gho et al., 1996; Olenchock et al., 2016). Thus, gradual adaptation to hypoxia can impart significant health advantages. This adaptation is partially mediated by the activation of hypoxia-inducible transcription factors (HIF) (Epstein et al., 2001; Ivan et al., 2001; Jaakkola et al., 2001), which promote erythropoiesis and angiogenesis to enable survival at lower oxygen levels (Haase, 2013; Pugh and Ratcliffe, 2003). While previous work has focused extensively on these transcriptional programs, metabolic adaptations to hypoxia remain incompletely understood at the whole-body level.

The metabolic responses to hypoxia are especially important given the striking observation that high-altitude residents are paradoxically protected against a range of metabolic conditions. More than 2 million individuals reside at an altitude greater than 4500m at an oxygen level that translates to an 11% fraction of inspired oxygen (F_i_O_2_) (compared to 21% at sea level) (Tremblay and Ainslie, 2021). Over a dozen studies have found that high-altitude residents have significantly lower rates of hyperglycemia, hypercholesterolemia, and obesity (Castillo et al., 2007; Díaz-Gutiérrez et al., 2016; Dünnwald et al., 2019; Koufakis et al., 2019; Lopez-Pascual et al., 2016, 2018; Merrill, 2020; Voss et al., 2014; Xu et al., 2017) and reduced mortality from coronary artery disease and stroke (Baibas et al., 2005; Faeh et al., 2009; Ortiz-Prado et al., 2021). Independent of altitude, oxygen deprivation alone can also have therapeutic benefits. For example, low blood hemoglobin levels are associated with improved glucose homeostasis, decreased insulin resistance, and lower LDL and triglyceride levels in mice and humans (Auvinen et al., 2021). Additionally, we recently showed that hypoxia exposure extends lifespan, reverses neurological lesions, and treats functional deficits in a mouse model of monogenic mitochondrial disease (Jain et al., 2016). These observations suggest that shifts in oxygen availability prompt a rewiring of organ-level and systemic metabolism that may be therapeutic.

Technological advances have enabled comprehensive, unbiased investigation of systemic metabolism, revealing the considerable metabolic flexibility and fuel rewiring of mammals in response to physiological stimuli. For example, endurance exercise in humans rapidly activates skeletal muscle glucose uptake and fatty acid oxidation (Lewis et al., 2010). Conversely, fasting activates a transition away from glucose utilization, mobilizing lipids, ketone bodies, and amino acids (Steinhauser et al., 2018). These metabolic switches optimize resource consumption to meet shifting energetic needs.

The findings of significant metabolic flexibility and sustained adaptation to hypoxia raise an important question—how do mammalian organs reorganize their fuel preferences and metabolic pathways to meet energy demands in oxygen-limited environments? The canonical view of hypoxia metabolism emphasizes a switch from oxidative phosphorylation to increased anaerobic glycolysis, suggesting a shift away from fatty acid metabolism towards glucose utilization. However, this perspective stems primarily from studies in cancer cell lines (Eales et al., 2016; Solaini et al., 2010; al Tameemi et al., 2019). Uniform whole-body reliance on anaerobic glycolysis is unlikely to be adaptive as it would result in rapid glucose depletion and severe lactic acidosis. Thus, it is more likely that different organs rewire their metabolism based on local oxygenation, organ-specific functions, and nutrient availability.

To test this hypothesis, we measured fuel uptake and metabolic flux in live mice exposed to acute and chronic ambient hypoxia using positron emission tomography and isotope-labeled fuel tracers. As expected, acute hypoxia increased glucose uptake in most organs while reducing the mitochondrial oxidation of glucose. Notable exceptions included skeletal muscle and brown fat, which serve as “glucose-saving” organs by decreasing glucose uptake in acute hypoxia. In chronic hypoxia, organs transformed their metabolism in unique ways. The heart increased its dependence on mitochondrial glucose oxidation, while the brain, kidney, and liver promoted fatty acid uptake to feed the TCA cycle. These organ-specific changes coincided with reduced body fat, dramatically reduced blood glucose levels, and improvements in hypoxia-induced perturbations in locomotor function. This comprehensive characterization elucidates how organs contribute differently to systemic hypoxia adaptation while producing a catabolic state. These findings highlight some mechanisms underlying hypoxia-induced improvements in metabolic fitness and organ resilience to oxygen deprivation.

## RESULTS

### Impaired locomotor function in acute hypoxia is rescued over time

To establish a model of hypoxia adaptation, we began by assessing spontaneous movement and fitness, since these traits are known to be acutely impaired in humans at high altitude. We housed young adult mice in three different oxygen tensions (21%, 11%, and 8% F_i_O_2_) for four different time periods (3 hours, 24 hours, 1 week, and 3 weeks) representing acute, subacute, and chronic hypoxia. 21% represents the oxygen tension of room air at sea level, and 11% and 8% correspond to altitudes of 4500m and greater than 5000m, respectively. These two F_i_O_2_s were chosen because they represent significant hypoxia that is survivable in mice and humans. Of note, 2 million people live above 4500m and 300,000 people live above 5000m (Tremblay and Ainslie, 2021).

To evaluate spontaneous movement, we conducted an open field assay, measuring distance travelled, speed, and resting time in a controlled atmosphere. Acutely, oxygen deprivation significantly reduced spontaneous movement and increased resting time, but by 3 weeks, mice exposed to hypoxia increased their movement to near normal levels **(Fig 1A-D)**. We also conducted pole tests as an integrated assessment of motor function, including coordination, strength, and speed (Balkaya and Endres, 2010). Mice were placed at the top of a pole facing up or down, and we measured the latency to descend. In both starting positions, mice exposed to 8% F_i_O_2_ for 24 hours took significantly longer to descend than mice exposed to 21% F_i_O_2_. By 3 weeks of exposure, however, the latency to descend among the hypoxia groups was comparable to the normoxic controls **(Fig 1E-F)**. Together, these results indicate that acute hypoxia reduces spontaneous motion and motor coordination, but adaptation over several weeks rescues these motor deficits.

**Figure 1.**
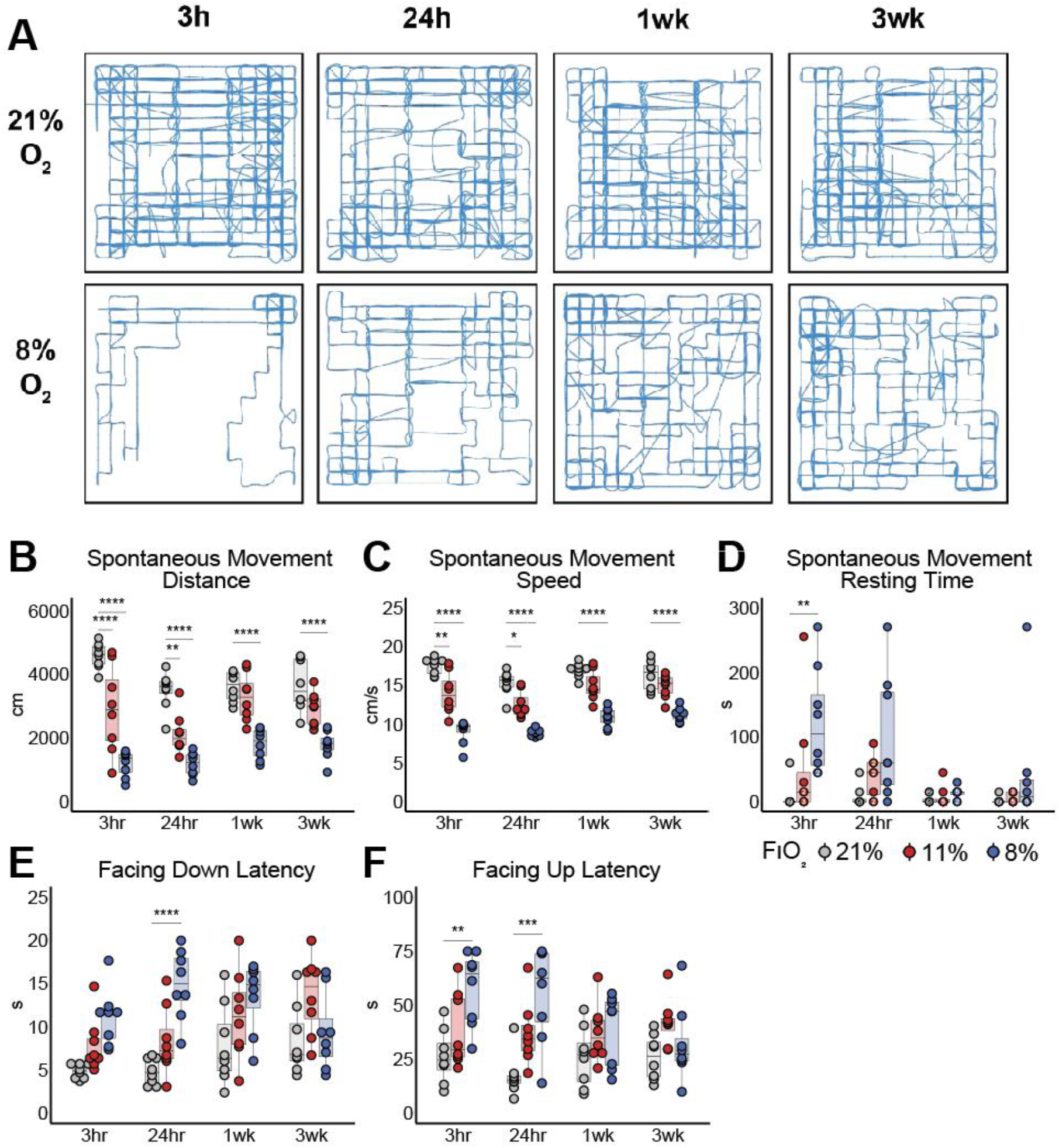
Acute hypoxia causes perturbations in spontaneous movement and motor coordination that attenuate over time. **(A)** Representative movement traces of male mice housed at 21% or 8% F_i_O_2_ and transferred to openfield chambers of matching F_i_O_2_. **(B-D) (B)** Total distance covered, **(C)** average speed, and **(D)** total resting time of mice placed in openfield chambers over 10 minutes. Mice were housed at 21% (grey), 11% (red), 8% (blue) F_i_O_2_ for 3 hours, 24 hours, 1 week, or 3 weeks. **(E-F)** Latency to descend (in seconds) down a pole after being placed at the top facing down **(E)** or facing up **(F)**. Statistics were calculated using two-way ANOVA and post-hoc Tukey correction. N = 8 biological replicates. *p<0.5, **p<.01, ***p<.001, ****p<.0001.

### Physiological responses to hypoxia including systemic metabolic rewiring

To identify the mechanisms by which acute locomotor defects improve under chronic hypoxia, we carefully monitored physiological markers known to respond to hypoxia, including hemoglobin, blood CO_2_, and body temperature. As expected, hemoglobin levels doubled after 3 weeks in 8% F_i_O_2_ **(Fig 2A)**, matching previous human studies (Lawrence et al., 1952; Pugh, 1964). Another well-known response to acute hypoxia is hyperventilation, which increases the expiration of CO_2_. As a result, acute hypoxia lowered total blood CO_2_, but over 3 weeks, this parameter returned to normal **(Fig 2B)**.

**Figure 2.**
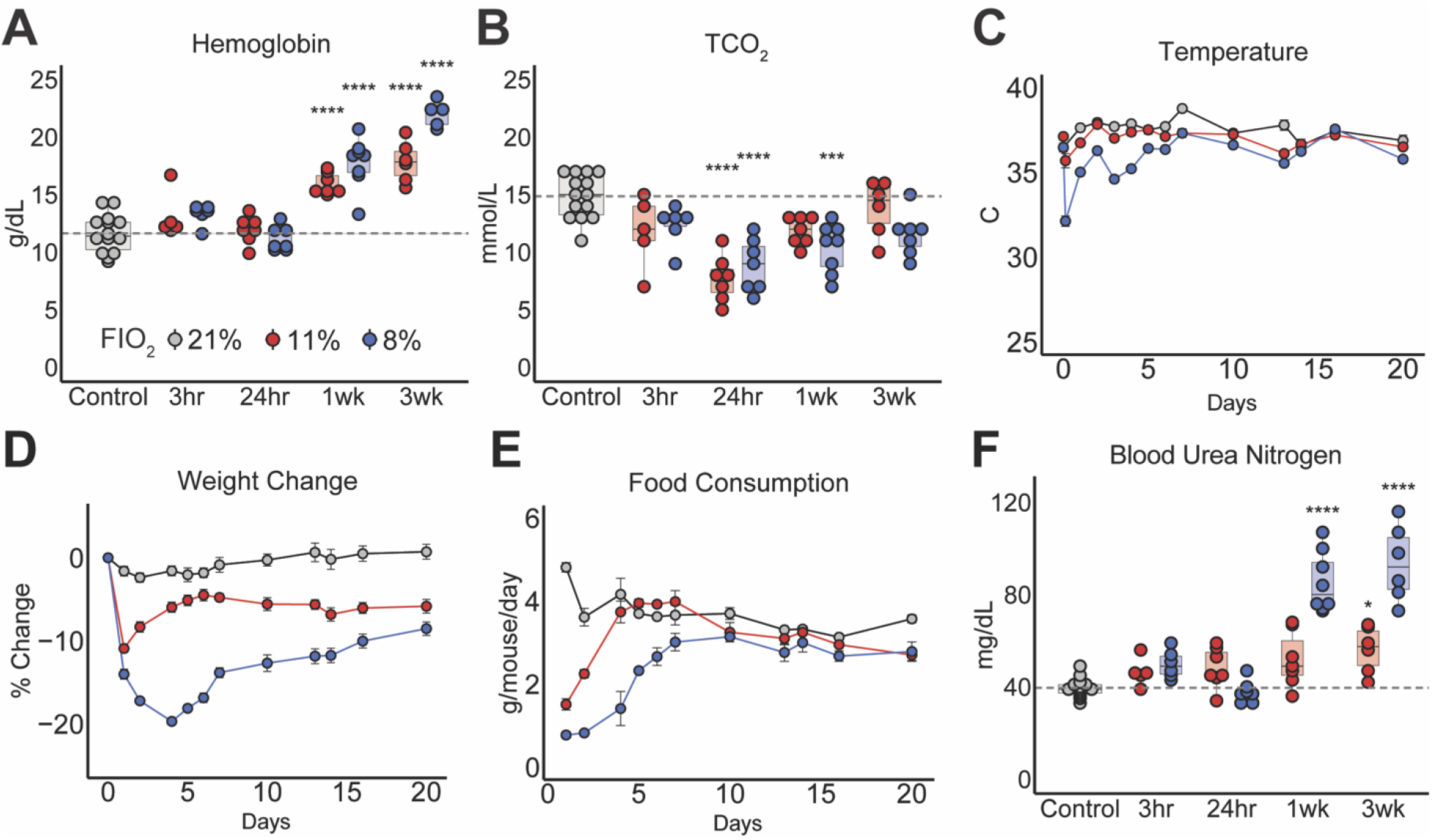
Physiological adaptation to hypoxia includes systemic metabolic rewiring. **(A-B) (A)** Blood hemoglobin concentrations and **(B)** Total CO_2_ measured from tail vein blood samples of mice housed in 21% (grey), 11% (red), or 8% F_i_O_2_ (blue) for 3 hours, 24 hours, 1 week, or 3 weeks. Reduced TCO_2_ levels indicate hyperventilation. **(C)** Highest detectable body temperatures measured with an infrared camera daily for one week and every 2-4 days for the following 2 weeks. Mean ± SEM are shown. For 11% F_i_O_2_ mice, body temperatures were significantly different from normoxic mice on days 7 and 13. For 8% F_i_O_2_ mice, body temperatures were significantly different from normoxic mice from 3 hours to 4 days and on days 6, 7, and 13. **(D)** Percent change in body weight from baseline. Mean ± SEM are shown. For 11% F_i_O_2_ mice, differences in body weight change were statistically significant from day 1 to 4 and from day 7 to 20 when compared to normoxic mice. For 8% F_i_O_2_ mice, differences in body weight change were statistically significant from day 1 to 20 when compared to normoxic mice. **(E)** Food consumption per mouse per day. N = 2 cages. Mean ± SEM are shown. For 11% F_i_O_2_ mice, food consumption was significantly different from normoxic mice on days 1 and 2. For 8% F_i_O_2_ mice, food consumption was significantly different from normoxic mice from day 1 to 5. **(F)** Blood urea nitrogen (BUN) concentrations measured from tail blood samples. Elevated BUN provides evidence of increased protein degradation. Statistics were calculated using two-way ANOVA and post-hoc Tukey correction. N = 8 biological replicates for all panels except (E). *p<0.5, **p<.01, ***p<.001, ****p<.0001.

In acute hypoxia, we also observed a dramatic drop in body temperature (anapyrexia), but this normalized over the following weeks **(Fig 2C)**. Hypoxia-induced anapyrexia has previously been observed in other mammals, including rats (Gautier et al., 1991), golden hamsters (Kuhnen et al., 1987), naked mole rats (Cheng et al., 2021), and humans (DiPasquale et al., 2015). The mechanism underlying this phenomenon remains unknown, but adaptive benefits in acute hypoxia may include preserved energy stores, increased hemoglobin-O_2_ binding affinity, and a dampened hyperventilation response to hypoxia (Wood, 1991). Indeed, lowering body temperature confers neuroprotection in hypoxic-ischemic encephalopathy (Reinboth et al., 2016). However, the reversal of hypoxia-induced anapyrexia over time indicates that persistently low body temperature is not required for survival in chronic hypoxia.

Given that thermogenesis requires significant energy expenditure, we asked whether mice exhibited changes in body weight and energy intake during their adaptation to hypoxia. Acute oxygen deprivation stimulated a significant decline in body weight that was partially reversed over time. Four days in 8% F_i_O_2_ lowered body weights by 20%, but by three weeks, weights had partially recovered and stabilized at 10% below baseline **(Fig 2D)**. The acute drop in body weight tracked closely with a similar reduction in food consumption **(Fig 2E)**. Previous studies have demonstrated that paired feeding rescues hypoxia-induced weight loss (Abu Eid et al., 2018), so depressed food intake is the likely cause of the observed weight loss. Research in humans has similarly demonstrated a rapid decline in food consumption at even more moderate hypoxia conditions (Shukla et al., 2005). We found that after one week of hypoxia, food consumption returned to normal, matching the time course of body temperature recovery.

The precise signaling cascade driving hypoxia-induced anorexia remains unknown. Increased leptin secretion, which promotes satiety, has been proposed as a possible mechanism, but human studies have produced conflicting results (Morishima and Goto, 2016; Shukla et al., 2005; Vats et al., 2004), perhaps linked to confounding variables such as ambient temperature, barometric pressure, and sleep at altitude. By contrast, our experimental setup allowed us to assess the effects of oxygen deprivation alone. We collected blood between 10am and 12pm without fasting mice to avoid influencing the results with exogenous interventions. From these hormone measurements, we found that leptin levels were elevated in mice housed at 8% F_i_O_2_ for 24 hours **(Fig 3A)**, which corresponds to the timepoint with the largest drop in food intake. In addition, levels of ghrelin, which promotes food intake, were significantly lower at 3 hours but returned to normal at 24 hours and were significantly elevated at 1 week **(Fig 3B)**. These data suggest that hormonal responses to hypoxia may contribute to acute anorexia.

**Figure 3.**
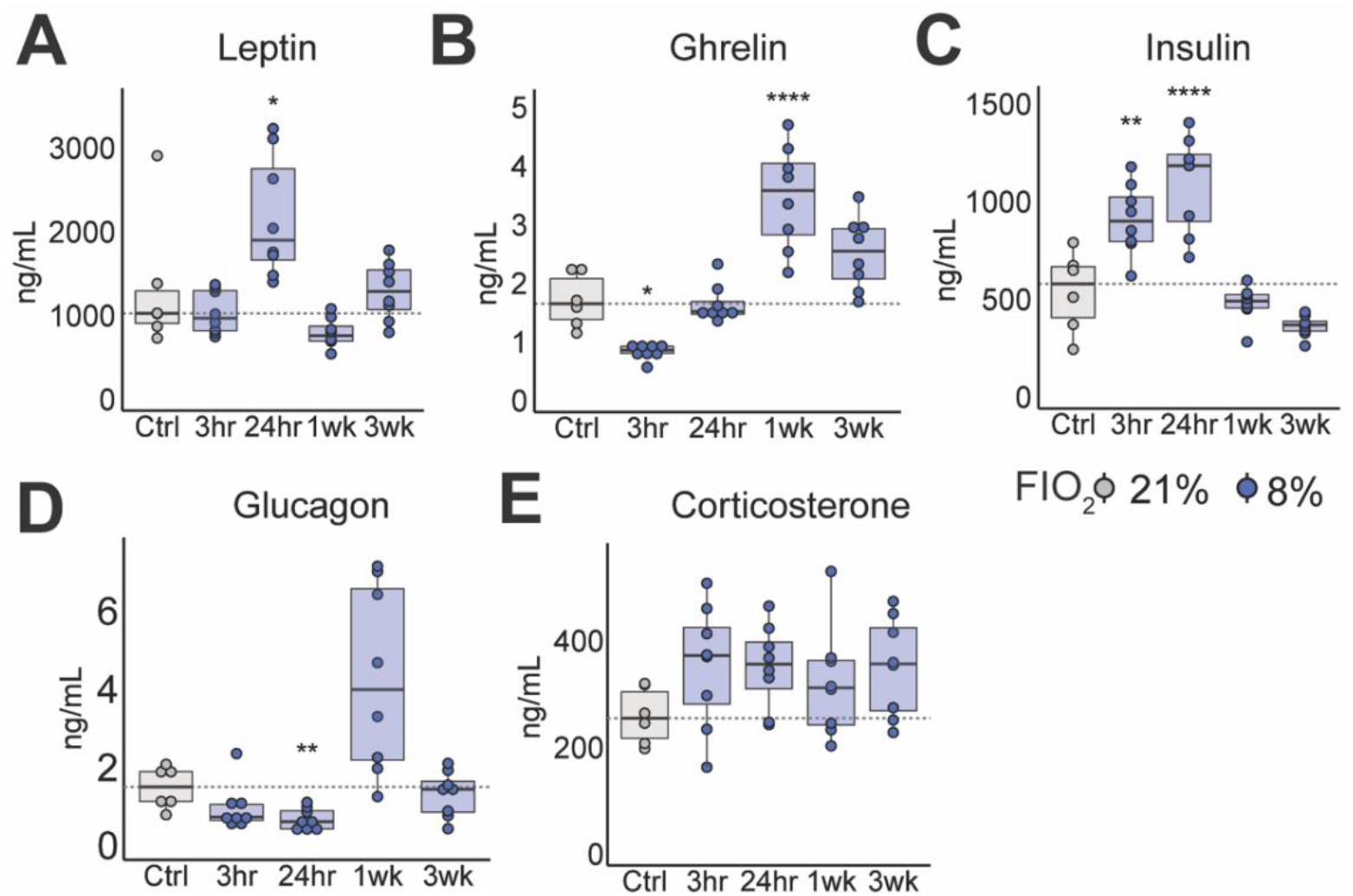
Hypoxia induces time-dependent changes in metabolic hormones. **(A-E)** Unfasted plasma levels of **(A)** leptin, **(B)** ghrelin, **(C)** insulin, **(D)** glucagon, and **(E)** corticosterone from tail blood samples after 3 hours, 24 hours, 1 week, and 3 weeks of hypoxia (8% F_i_O_2_) treatment (blue). Controls (grey) were housed in 21% F_i_O_2_. Statistics were calculated using one-way ANOVA and post-hoc Tukey correction. N = 6-8 biological replicates. *p<0.5, **p<.01, ***p<.001, ****p<.0001.

Regardless of its cause, the rapid and dramatic weight loss in hypoxia led us to hypothesize that oxygen-deprived mice may activate metabolic starvation-response pathways. Consistent with this hypothesis, hypoxia produced a significant increase in blood urea nitrogen (BUN) that was time- and oxygen dose-responsive **(Fig 2F)**. Blood urea is produced from ammonia generated by protein degradation, and this urea is excreted by the kidney. Therefore, elevated BUN can indicate kidney injury or elevated rates of protein catabolism (Giesecke et al., 1989). However, we observed that serum creatinine levels, another marker of kidney injury, remained below the detectable threshold throughout hypoxia treatment. Therefore, our observation of elevated BUN indicates a perturbed nitrogen balance most likely caused by protein catabolism. Consistent with these findings, altitude studies have found that chronic hypoxia reduces skeletal muscle mass (Hoppeler et al., 1990; MacDougall et al., 1991), and oxygen deprivation in cell lines activates protein catabolism to meet energy demands (Frezza et al., 2011). Together, our findings highlight that adaptation to hypoxia involves systemic metabolic rewiring, lowering energy intake and thermogenesis acutely and promoting chronic weight loss and protein breakdown.

### Chronic hypoxia lowers blood glucose levels and alters organ-level glucose uptake

Given the dramatic metabolic changes in hypoxia and the prior evidence of reduced diabetes incidence among high-altitude populations, we asked whether hypoxia alone could alter systemic glucose homeostasis. Exposure to 8% F_i_O_2_ induced a significant time- and dose-dependent reduction in fasted and fed glucose levels **(Fig 4A)**. As expected, normoxic mice had fasted glucose levels around 100 mg/dL and fed glucose levels around 150 mg/dL. At 3 hours of treatment, fasted and fed glucose levels were slightly lower before recovering to normal at 24 hours. By 1 week, though, both fed and fasted blood glucose levels cratered to ~55 mg/dL. Despite the return to normal food consumption at 3 weeks, fasted and fed blood glucose levels were both less than 40 mg/dL. This dramatic 3-4-fold decrease in circulating glucose was not accompanied by any overt detrimental effects. To the contrary, behavior recovered to normoxic states at chronic timepoints **(Fig 1)**.

**Figure 4.**
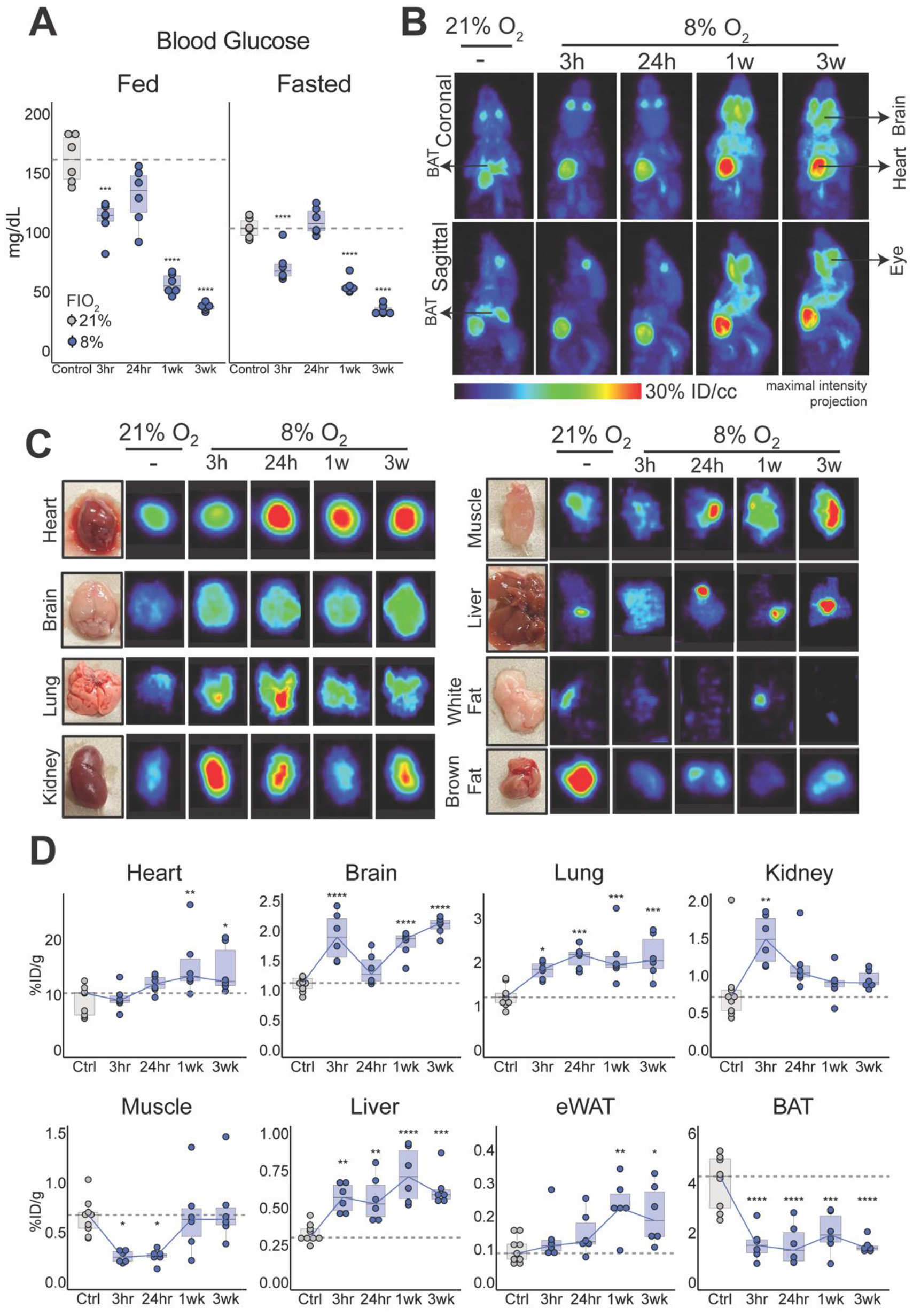
Chronic hypoxia causes hypoglycemia and alters organ-specific glucose uptake. **(A)** Fed and fasted blood glucose measurements after 3 hours, 24 hours, 1 week, and 3 weeks of hypoxia (8% F_i_O_2_) treatment (blue). Controls (grey) were housed in 21% F_i_O_2_. **(B)** Representative coronal and sagittal images of mouse PET scans conducted 30 minutes after tail vein injection of the glucose analogue 2-Deoxy-2-[^18^F]fluoro-D-glucose (FDG). **(C)** Representative PET scan images of organs extracted from mice 60 minutes after tail vein injection of FDG. Maximum values (% ID/cc) vary per organ: Heart: 40, Brain: 20, Lung: 10, Kidney: 5, Muscle: 7, Liver: 7, White Fat: 2, Brown Fat: 10. **(D)** Radioactive signal from each organ after extraction as measured by a gamma counter. Values were decay-corrected based on the time of FDG injection and the time of measurement. Statistics were calculated using one-way ANOVA and post-hoc Tukey correction. N = 6-9 biological replicates. *p<0.5, **p<.01, ***p<.001, ****p<.0001. eWAT: epidydimal white adipose tissue, BAT: brown adipose tissue.

To explain this observation, we considered hypoxia-induced changes in pancreatic islet endocrine activity. Insulin levels were elevated at acute timepoints but returned to normal at 1 week and 3 weeks in 8% F_i_O_2_ **(Fig 3C)**. Conversely, glucagon levels were decreased at 24 hours and varied greatly at 1 week before returning to normal **(Fig 3D)**. Neither hormone was significantly perturbed in chronic hypoxia. Therefore, changes in insulin and glucagon production are unlikely to explain the dramatic hypoglycemia in chronic hypoxia.

We therefore asked whether organs increased glucose consumption as a function of time in hypoxia. We measured organ-specific glucose uptake using PET-CT imaging with an injected 2-deoxy-2-[^18^F]fluoro-D-glucose (FDG) tracer. FDG is taken up by cells through glucose transporters; after phosphorylation, the molecule is trapped within cells and does not undergo further metabolism (Gallagher et al., 1978). Thus, the radioactive ^18^F signal can be imaged as a measure of glucose uptake. We conducted FDG PET scans in mice housed at 21% and 8% F_i_O_2_ for 3 hours, 24 hours, 1 week, and 3 weeks. We also extracted organs and used a gamma counter to quantify the ^18^F signal from each organ (Zhao et al., 2021).

In normoxia, the major glucose consumers were the heart and brown adipose tissue (BAT), followed by the lungs and brain **(Fig 4B)**. In response to hypoxia, most organs acutely increased their glucose uptake, but the kinetics differed across organs **(Fig 4C-D, S1A-B)**. For example, glucose uptake increased after as little as 3 hours in hypoxia for the brain and kidney, compared to 1 week for the heart. These organ-specific differences may implicate the time required for organs to adapt to ischemic injury or systemic hypoxic events.

These results are consistent with the classical teachings of hypoxia adaptation in proliferating cells. In this model, hypoxia promotes glucose uptake, upregulates glycolytic enzymes, and increases lactate excretion as the end-product of anaerobic glycolysis (al Tameemi et al., 2019). Contrary to this traditional view and the overall trend of increased glucose uptake, two organs substantially decreased their glucose consumption in acute hypoxia: skeletal muscle and BAT. In skeletal muscle, glucose uptake returned to baseline at 3 weeks, a pattern that mirrors the acute impairment and subsequent improvement in locomotor activity. Reduced movement during acute hypoxia likely preserves circulating glucose for other organs because skeletal muscle contraction promotes glucose uptake (Ryder et al., 2001).

In BAT, glucose uptake was decreased by nearly five-fold persistently across acute and chronic hypoxia. Interestingly, both organs contribute to thermoregulation; skeletal muscle activity enables shivering thermogenesis, and BAT metabolism drives non-shivering thermogenesis (Betz and Enerbäck, 2018). The acute suppression of glucose uptake in both organs coincided with a significant drop in body temperature **(Fig 2C)**. A study in naked mole rats showed that exposure to 7% F_i_O_2_ rapidly lowered body temperature and reduced UCP1 expression in brown fat, which is required for thermogenesis (Cheng et al., 2021). Therefore, the improvement in body temperature over time may emerge due to a switch away from BAT thermogenesis as skeletal muscle metabolism recovers.

### Chronic hypoxia reduces fat accumulation and alters organ-level uptake of free fatty acids

After observing such dramatic changes in glucose utilization, we asked whether hypoxia affects additional major fuel sources. To this end, we characterized hypoxia-induced changes in systemic and organ-specific fatty acid metabolism. We measured the fat content in mice by conducting dual-energy X-ray absorptiometry (DEXA) scans and weighing epidydimal white adipose tissue (eWAT), a visceral fat depot. Chronic hypoxia reduced total fat mass and the weight of eWAT depots **(Fig 5A)**. Chronic hypoxia also reduced total lean mass, but to a smaller degree than the reduction in fat mass **(Fig S2A-C)**. Strikingly, we observed reduced adiposity despite a recovery in food intake after the first week of hypoxia treatment. Along with the observed reduction in blood glucose levels, these data provide experimental support to the hypothesis that chronic adaptation to hypoxia can improve metabolic health by increasing catabolism of stored fuels.

**Figure 5.**
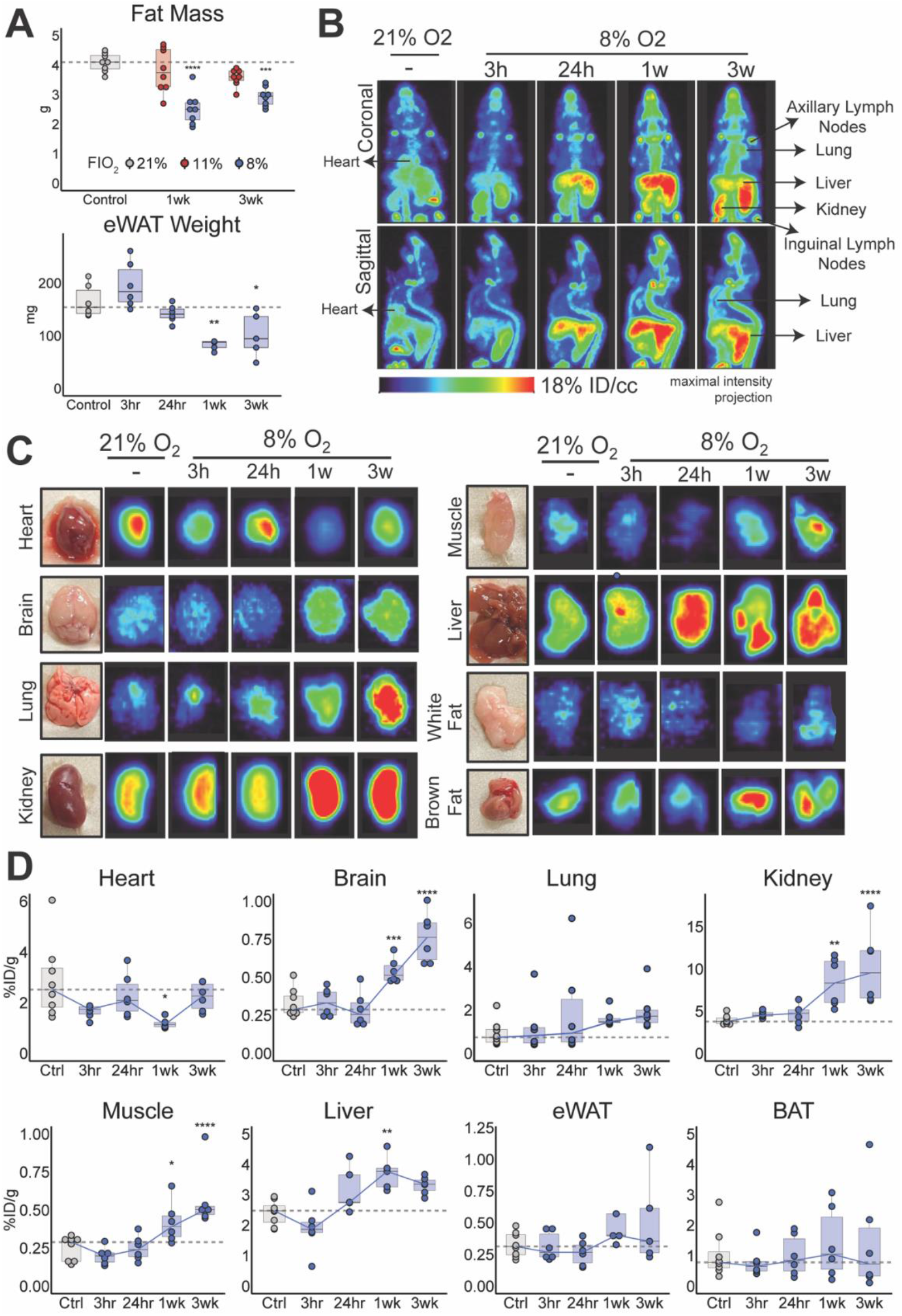
Chronic hypoxia reduces lipid accumulation and alters organ-specific free fatty acid uptake. **(A)** Fat mass measured by DEXA scan and weights of eWAT depots in mice housed at 21% (grey), 11% (red), and 8% (blue) F_i_O_2_. **(B)** Representative coronal and sagittal images of mouse PET scans conducted 30 minutes after tail vein injection of the palmitate analogue 18-[18F]fluoro-4-thia-palmitate (FTP). **(C)** Representative PET scan images of organs extracted from mice 60 minutes after tail vein injection of FTP. Maximum values (% ID/cc) vary per organ: Heart: 20, Brain: 20, Lung: 20, Kidney: 50, Muscle: 7, Liver: 65, White Fat: 2, Brown Fat: 15. **(D)** Radioactive signal from each organ after extraction as measured by a gamma counter. Values were decay-corrected based on the time of FDG injection and the time of measurement. Statistics were calculated using one-way ANOVA and post-hoc Tukey correction. N = 4-8 biological replicates. *p<0.5, **p<.01, ***p<.001, ****p<.0001. eWAT: epidydimal white adipose tissue, BAT: brown adipose tissue.

To investigate the mechanism underlying reduced fat accumulation in chronic hypoxia, we measured fatty acid uptake in organs. Analogous to the FDG PET scans, we conducted PET scans using 18-[18F]fluoro-4-thia-palmitate (FTP), a metabolically trapped free fatty acid (FFA) tracer (DeGrado et al., 2000). FTP resembles the saturated fatty acid palmitate, the most abundant FFA in circulation. FTP undergoes mitochondrial uptake through the carnitine shuttle before being trapped in the mitochondria (DeGrado et al., 2006).

As expected, FTP PET scans showed that in normoxia, the heart, liver, and kidney were the most significant consumers of circulating palmitate **(Fig 5B)**. On the other hand, the brain exhibited ten-fold lower FFA uptake than the heart. Acute hypoxia had no significant effect on FFA metabolism across organs. Meanwhile, chronic hypoxia prompted a significant increase in FFA uptake in the liver, kidney, brain, muscle, and spleen **(Fig 5C-D, S2E)**. These unexpected findings contradict classical teachings from cancer cell lines of decreased fatty acid uptake and oxidation in hypoxia (Du et al., 2017; Liu et al., 2014).

### Hypoxia rewires metabolic flux of major fuel sources into the TCA cycle

Given the importance of oxygen in mitochondrial oxidative metabolism, we next investigated how hypoxia affects the shuttling of circulating fuels into the TCA cycle. We conducted separate bolus injections of the stable isotopes U-^13^C-glucose and U-^13^C-palmitate and extracted organs 20 minutes later. The concentrations of these tracers in plasma varied slightly across conditions, reflecting different baseline concentrations of circulating glucose and palmitate in hypoxia **(Fig S4A)**. Next, we traced the conversion of circulating carbons from glucose and palmitate into the TCA intermediates succinate, fumarate, and malate and the TCA-derived amino acid glutamate. The fractional labeling of TCA metabolites, as determined by LC-MS analysis of extracted organs, represents the use of the injected fuels to feed the TCA cycle **(Fig 6A, S3B)**. Because the tracer measurements were not conducted at steady state, the exact fractional contributions of glucose and palmitate to downstream metabolites cannot be determined, and enrichments of the isotope label cannot be precisely compared across different organs. Nevertheless, increased enrichment of the isotope label in TCA metabolite implies increased production of TCA intermediates from the labeled fuel relative to other sources.

**Figure 6.**
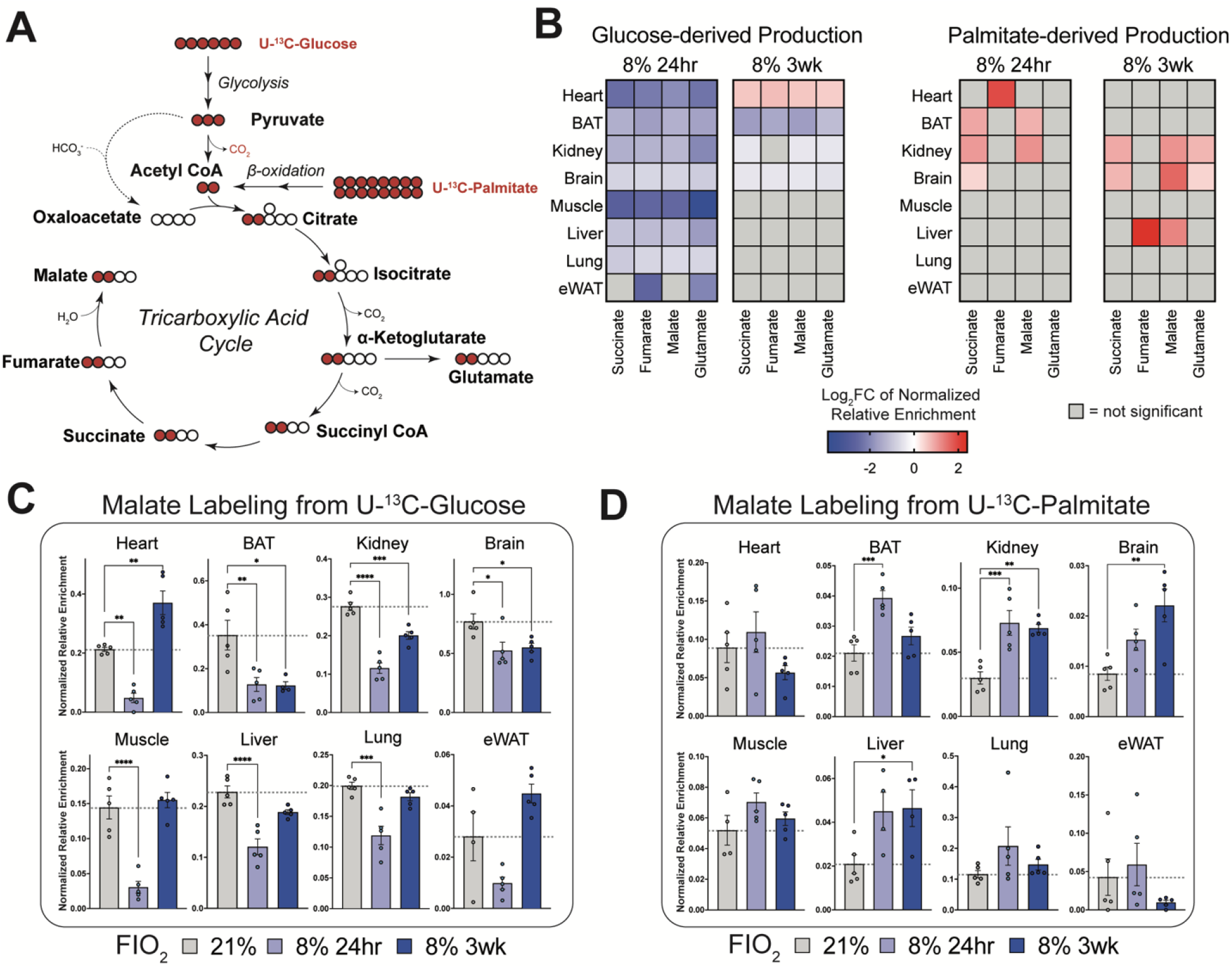
Hypoxia rewires metabolic flux of major fuel sources into the TCA cycle. **(A)** Schematic showing entry of carbons from glucose and palmitate to the TCA cycle. Red circles represent ^13^C-labeled carbons. Labeling patterns correspond to ^13^C-Acetyl CoA passing through one round of the TCA cycle. **(B)** Heatmap showing the log fold change in normalized relative enrichment of the isotope label compared to the 21% F_i_O_2_ condition. Normalized relative enrichment was calculated by determining the fraction of carbons in TCA metabolites that were ^13^C-labeled and subsequently normalizing to the ^13^C enrichment of the fuel (glucose or palmitate) in plasma. Greater normalized relative enrichment indicates increased production of the target metabolite from the injected tracer compared to other sources. Only statistically significant changes are shown. **(C-D)** Normalized relative enrichment of malate 20 minutes after injection with **(C)** ^13^C-glucose or **(D)** ^13^C-palmitate. Mean ± SEM are shown. Statistics were calculated using one-way ANOVA and post-hoc Dunnett’s test for multiple comparisons to a control group. N=4-5 biological replicates. *p<0.5, **p<.01, ***p<.001, ****p<.0001.

As expected, we found that in every organ, acute hypoxia reduced the production of TCA metabolites from circulating glucose **(Fig 6C, S5A)**. This finding was most pronounced in the heart and skeletal muscle **(Fig 6B)**. This pattern coincided with increased glucose uptake in many organs, suggesting the consumption of glucose for anaerobic glycolysis or other, non-TCA fates. This finding is consistent with a hypoxia-induced blockade of pyruvate dehydrogenase (PDH) activity by HIF (Papandreou et al., 2006). Notably, we observed no such blockade of mitochondrial fatty acid oxidation in acute hypoxia. In all organs, relative palmitate-derived production of TCA metabolites remained steady or even elevated in acute hypoxia **(Fig 6B, 6D, S5B)**.

In chronic hypoxia, however, organs exhibited distinct patterns of glucose oxidation into the TCA cycle. In the heart, glucose-derived production of TCA metabolites was significantly increased relative to normoxia. On the other hand, brown fat, the kidney, and the brain exhibited a significant decrease in glucose-derived TCA metabolites relative to normoxia **(Fig 6C, S5A)**.

Meanwhile, chronic hypoxia significantly increased the contribution of circulating palmitate to the TCA cycle in the kidney, brain, and liver **(Fig 6D, S5B)**. Of note, this 2-3x increase in fatty acid oxidation into the TCA cycle mirrored the 2-3x increase in FTP-uptake observed in these organs during chronic hypoxia. This upregulation of fatty acid uptake and oxidation in chronic hypoxia belies classical teachings of suppressed oxidative metabolism during hypoxia.

Based on these observations, we identify distinct metabolic classes of organs in chronic hypoxia. The heart stands alone as an increased glucose oxidizer, shuttling carbons from circulating glucose into the mitochondria to feed the TCA cycle. On the other hand, the kidney, liver, and brain mobilize circulating palmitate for mitochondrial metabolism. Finally, BAT suppresses the entry of glucose-derived carbons into the TCA cycle. Altogether, these findings highlight the unique metabolic contributions of different organs to systemic hypoxia adaptation.

## DISCUSSION

Acute disruptions in oxygen availability can be lethal. Nevertheless, adaptation to hypoxia at high altitude confers significant metabolic benefits, including improved glycemic control (Castillo et al., 2007), reduced coronary artery disease mortality (Baibas et al., 2005), and improved exercise capacity (Matheson et al., 1991). Mammals can adapt to hypoxia through several known mechanisms, including erythropoiesis (Haase, 2013; Pugh and Ratcliffe, 2003), angiogenesis (Pugh and Ratcliffe, 2003; Vilar et al., 2008), and shifts in renal bicarbonate handling (Zouboules et al., 2018). However, the study of metabolic adaptations to hypoxia has thus far been largely limited to the study of cancer cell lines. As a result, the canonical view of hypoxia metabolism focuses on a cellular shift from oxidative phosphorylation to glycolytic energy production. However, hypoxic metabolism at the organ and whole-body resolutions has not been systematically characterized.

Here, we show that adaptation to hypoxia entails a systemic metabolic transformation, lowering circulating blood glucose and diminishing body weight and adiposity. We found that these metabolic effects were not caused by persistent endocrine changes. Rather, this whole-body metabolic phenotype was accompanied by organ-specific shifts in the uptake and mitochondrial oxidation of circulating fuel sources.

Acutely, most organs responded to hypoxia by increasing glucose uptake while limiting the entry of glucose-derived carbons into the TCA cycle. This pattern indicates increased usage of glucose for non-mitochondrial metabolism (including anaerobic glycolysis), which matches the classical view of adaptation to hypoxia. Meanwhile, brown fat and skeletal muscle deviated from this trend by decreasing their glucose uptake in acute hypoxia, serving as “glucose savers” for other organs that may have a greater need for glucose. Notably, the functions of both organs (thermogenesis and locomotion, respectively) were impaired in acute hypoxia. The partitioning of glucose in acute hypoxia implies that the non-vital functions of thermogenesis and locomotion are compromised to prioritize vital functions of other organs.

In chronic hypoxia, metabolic reprogramming varied significantly across organs **(Fig 7)**. The heart consumed significant amounts of glucose and fatty acids in normoxia but in chronic hypoxia, shifted toward glucose uptake to feed the TCA cycle. These findings are consistent with prior evidence that chronic hypoxia increases the heart’s dependence on glucose (Liu et al., 2021). Meanwhile, the brain shifted in the opposite direction, relying more heavily on circulating fatty acids to feed the TCA cycle. Despite this shift in mitochondrial metabolism, the brain still increased its glucose uptake significantly, suggesting alternate, non-mitochondrial fates for this glucose. The liver and kidney, which were both major fatty acid consumers in normoxia, further increased their fatty acid oxidation in hypoxia by 2-3-fold. Finally, in BAT, glucose uptake and oxidation remained suppressed in chronic hypoxia, and skeletal muscle returned both glucose uptake and oxidation to normal levels.

**Figure 7.**
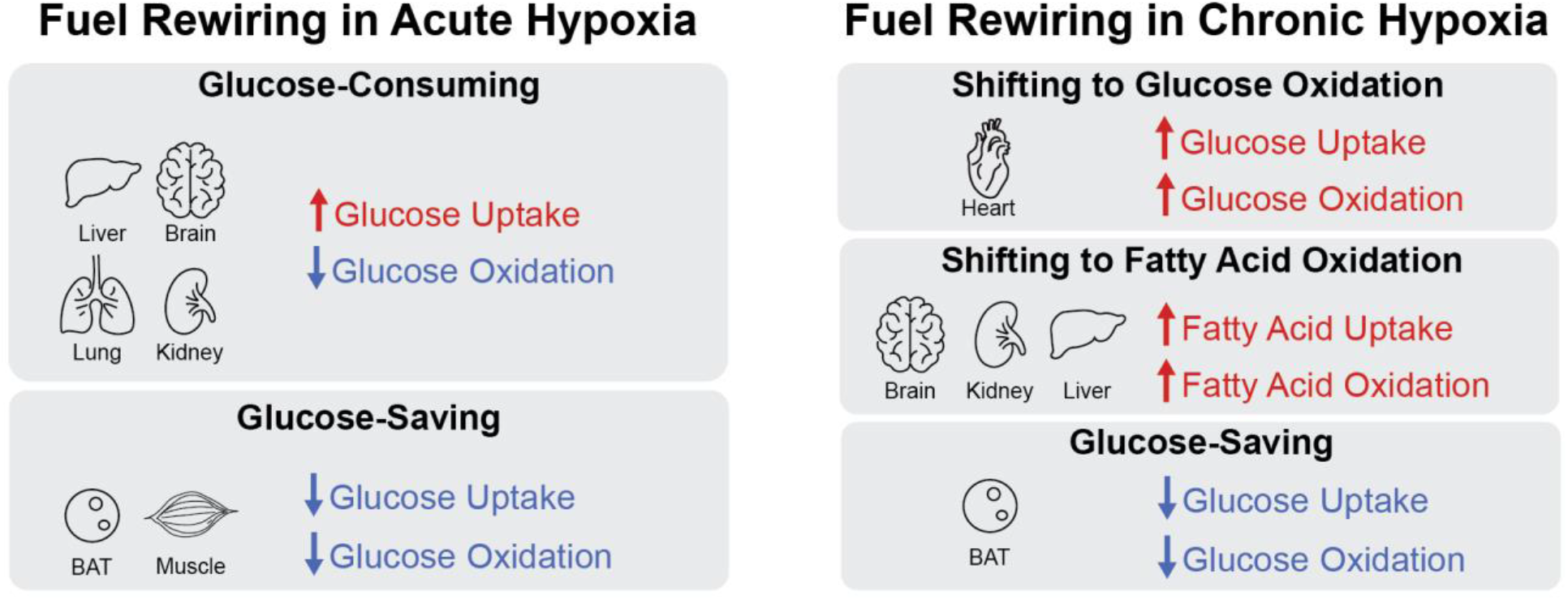
Summary of organ-specific fuel rewiring in physiological adaptation to hypoxia. Classification of organs that exhibit significant alterations in uptake and mitochondrial oxidation of the circulating fuel sources glucose and palmitate in acute and chronic hypoxia.

Our findings support different rationales for metabolic rewiring in acute versus chronic hypoxia. In acute hypoxia, the primary goal may be to conserve fuel sources for organs with the most vital functions (e.g., heart, brain, liver) at the expense of more acutely dispensable functions (e.g., locomotion or precise thermoregulation). In this case, suppressing energy demand in such organs preserves both oxygen and fuel supply. However, on the chronic timescales, organisms likely need to reactivate normal physiological functions, requiring efficient ATP production across all organs. This is achieved by selectively partitioning glucose and fatty acids depending on energy demands, metabolic flexibility, and local oxygenation of different organs.

Considering the increased oxygen demands of fatty acid oxidation, we did not expect to identify organs that significantly promoted fatty acid oxidation in chronic hypoxia (Kessler and Friedman, 1998). However, different organs have different baseline pO_2_ levels and oxygen-consuming needs in normoxia. Well-perfused organs may therefore maintain their fatty acid oxidation in hypoxia, thus saving glucose for less well-perfused locations that may rely on glucose uptake. Our data indicate that this phenomenon is maintained by an organ-specific increase in fatty acid oxidation in the liver, kidney, and brain.

The identification of the brain as an increased fatty acid consumer in chronic hypoxia was particularly unexpected. Ordinarily, the brain relies on glucose consumption for energy supply and, when fasted, on ketone bodies (Owen et al., 1967). Nevertheless, recent studies have demonstrated a capacity for low rates of fatty acid oxidation in the brain (White et al., 2020). The brain’s increased fatty acid oxidation in chronic hypoxia suggests an improved supply of oxygen over the course of adaptation to hypoxia. Indeed, severe chronic hypoxia has been shown to promote angiogenesis in the brain (Patt et al., 1997). Moreover, the hypoglycemia in chronic hypoxia may activate molecular starvation signals, which promote glial fatty acid oxidation and ketogenesis to supply fuel to neurons (Silva et al., 2022). In this study, we focused on organ-level metabolic rewiring. Future studies will focus on specific cell types and regional effects within organs.

Additionally, our current work was conducted on a normal chow diet. It is possible that dietary manipulations (e.g., high fat diet, ketogenic diet, or amino acid-restricted diets) will result in unique whole-body rewiring in hypoxia. Interestingly, an injection of ketone bodies has previously been shown to extend lifespan in severe, acute hypoxia (Rising and D’Alecy, 1989). Future work will incorporate these diets, trace additional fuel sources (lactose, ketones, and amino acids), and identify other metabolic fates of circulating nutrients (pentose phosphate pathway, gluconeogenesis, and one-carbon metabolism).

Collectively, our findings carry relevance for many metabolic conditions. In chronic hypoxia, we observed an increase in the consumption of circulating fuels by most organs. This may drive a starvation-like program that prevents cardiometabolic diseases caused by excess nutrient storage. Indeed, non-alcoholic fatty liver disease, type 2 diabetes, and atherosclerosis are all worsened by weight gain, hyperglycemia, and lipid accumulation. In chronic hypoxia, these risk factors were all reversed (Fig 2D, 4A, 5A), which may contribute to the lower burden of chronic metabolic diseases among high-altitude populations. Furthermore, organ-specific fuel rewiring in adaptation to hypoxia offers a metabolic starting point for maintaining function during acute disruptions in oxygen supply. Altogether, our results highlight the remarkable mammalian capacity for metabolic flexibility during hypoxia, which we could eventually harness for therapeutic ends.

## ACKNOWLEDGEMENTS

We thank all members of the Jain lab and Francoise Chanut for thoughtful discussions and review of the manuscript. We thank the UC Davis Mouse Metabolic Phenotyping Core for mouse hormone concentration measurements. We thank Ryan Tang for his technical assistance with PET/CT scans and associated radiotracer injections. We acknowledge the UCSF Radiopharmaceutical Facility for providing the FDG and [^18^F]fluoride ion for subsequent preparation of FTP. ADM was supported by the National Institute of General Medical Sciences (NIGMS) Medical Scientist Training Program, Grant T32GM141323. IHJ was supported by NIH DP5OD026398. IHJ, BBQ and AMB were supported by Defense Advanced Research Projects Agency, Biological Technologies Office (BTO) Program: Panacea issued by DARPA/CMO under Cooperative Agreement No. HR0011-19-2-0018

## AUTHOR CONTRIBUTIONS

IHJ, ADM, YZ, BBQ, AMB conceived the project and performed the experiments. IHJ, ADM, YZ designed the experiments and analyzed the data. ADM, IHJ wrote the manuscript. COYF, JEB prepared PET radiotracers. HV, YS designed PET radiotracers and assisted with data analysis.

## SUPPLEMENTAL FIGURES

**Figure S1.**
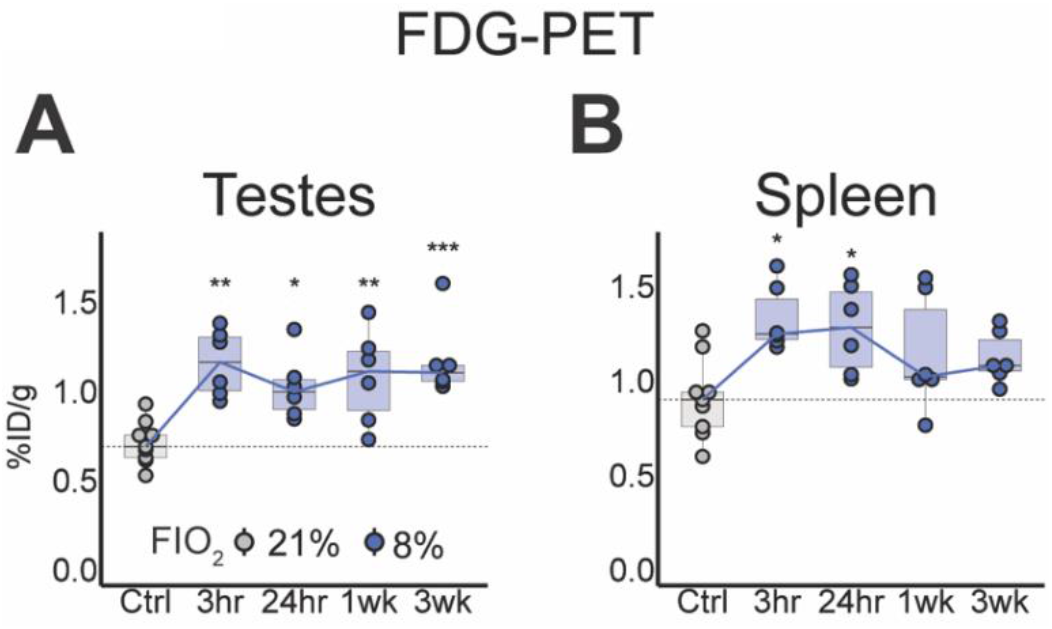
FDG-PET quantification in testes and spleen. **(A-B)** Radioactive signal from **(A)** testes and **(B)** spleen after extraction as measured by a gamma counter. Values were decay-corrected based on the time of FDG injection and the time of measurement. Statistics were calculated using one-way ANOVA and post-hoc Tukey correction. N = 5-6. *p<0.5, **p<.01, ***p<.001, ****p<.0001.

**Figure S2.**
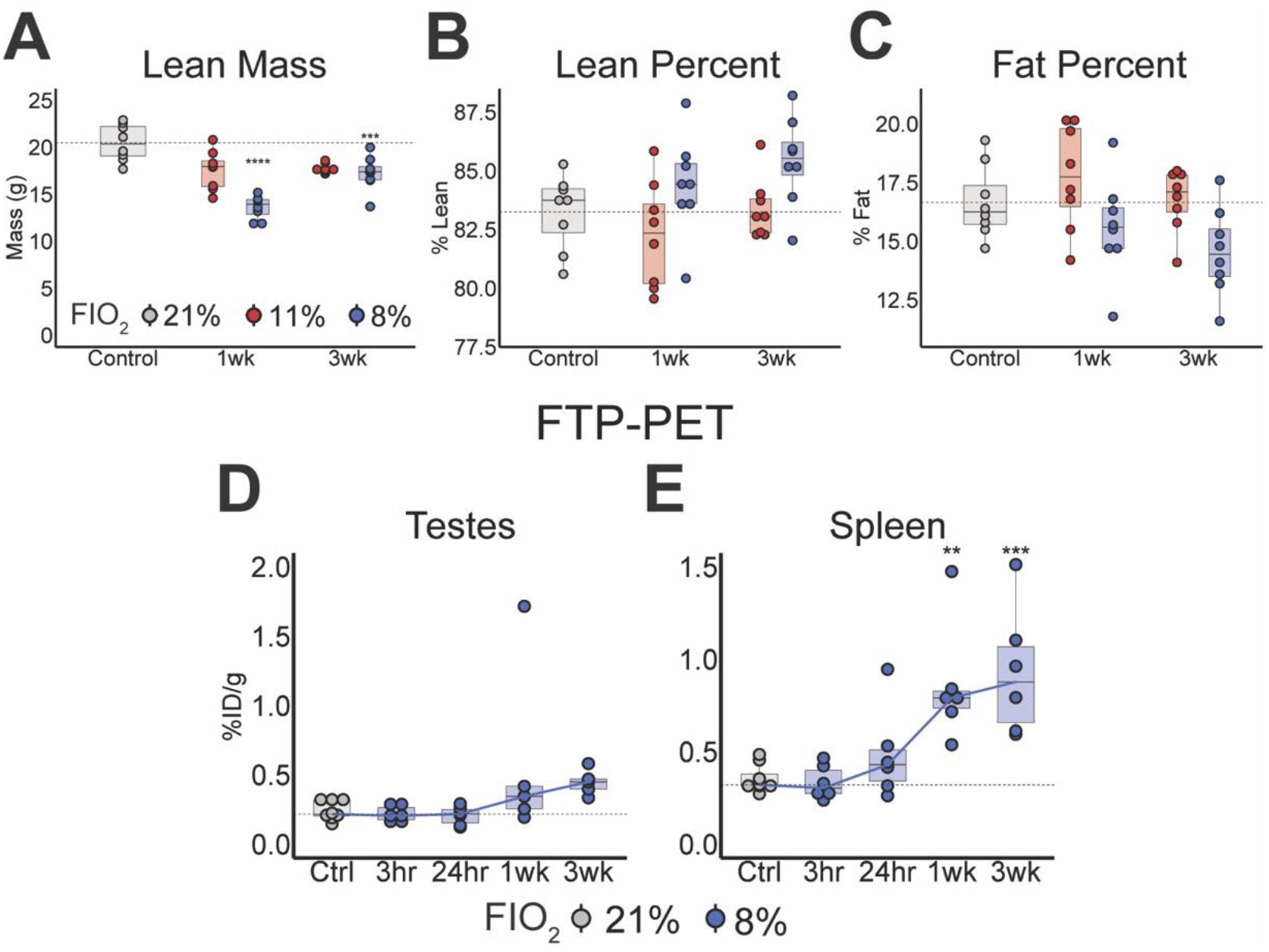
Whole-body lean and fat content and FTP-PET signal in testes and spleen. **(A-C)** DEXA scan measurements of **(A)** lean mass, **(B)** lean percent, **(C)** fat percent. **(D-E)** Radioactive signal from **(D)** testes and **(E)** spleen after extraction as measured by a gamma counter. Values were decay-corrected based on the time of FTP injection and the time of measurement. For panels (A) to (C), statistics were calculated using two-way ANOVA and post-hoc Tukey correction. For panels (D) to (F), statistics were calculated using two-way ANOVA and post-hoc Tukey correction. N = 7 for panels (A) to (C), and N = 5-6 for panels (D) and (E). *p<0.5, **p<.01, ***p<.001, ****p<.0001.

**Figure S3.**
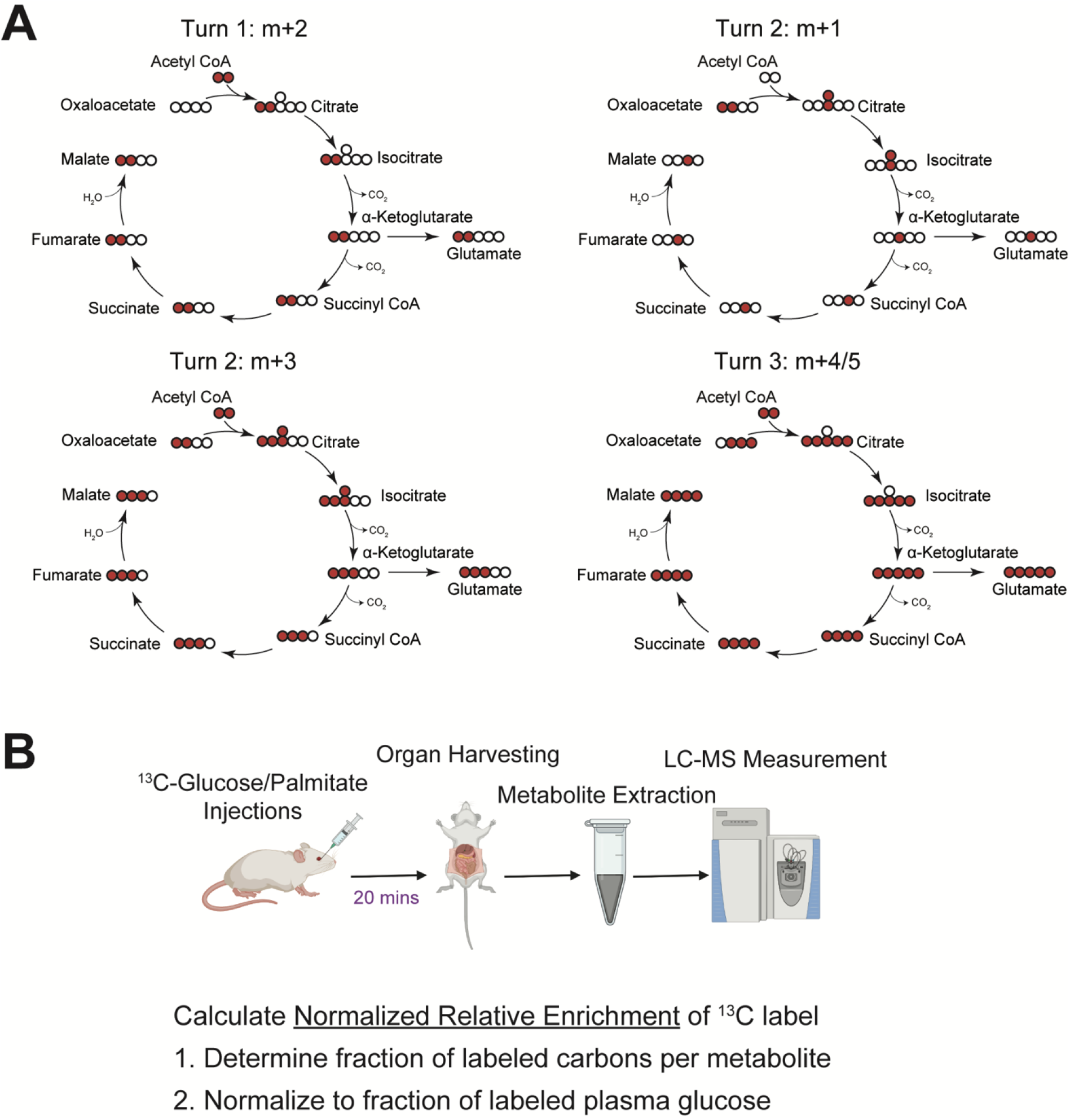
Schematic of isotope-labeled tracer experiments. **(A)** Depiction of possible oxidative pathways for the generation of different isotopomers of TCA metabolites (Inigo et al., 2021). All pathways involve at least one fully labeled acetyl CoA molecule entering the TCA cycle. **(B)** Schematic showing steps of isotope-labeled tracer experiments and calculation of normalized relative enrichment values. Created with BioRender.com.

**Figure S4.**
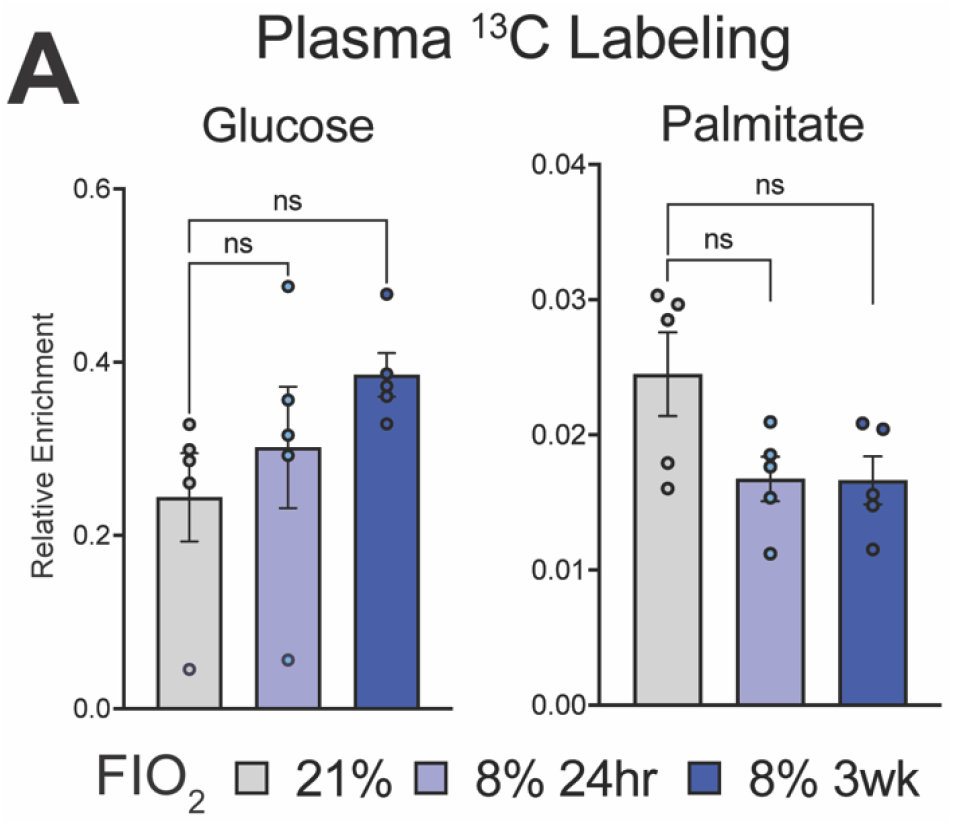
Plasma labeling of tracers. **(A)** Plasma enrichment of ^13^C isotope label 20 minutes after injection with either U-^13^C-glucose or U-^13^C-palmitate. Mean ± SEM are shown. Statistics were calculated using one-way ANOVA and post-hoc Dunnett’s test for multiple comparisons to a control group. N=5. *p<0.5, **p<.01, ***p<.001, ****p<.0001.

**Figure S5.**
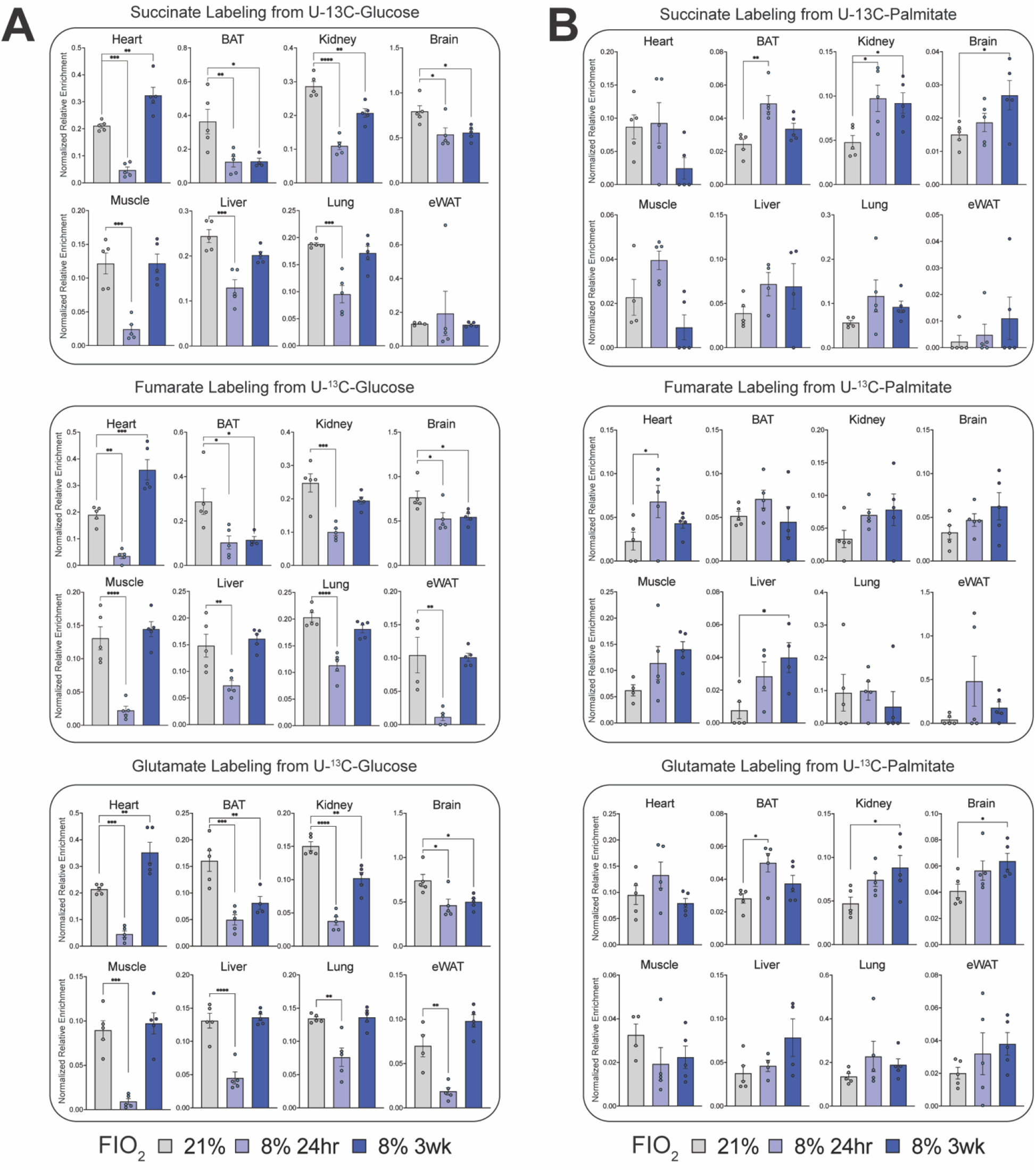
Labeling of TCA metabolites in organs. **(A-B)** Normalized relative enrichment of the ^13^C isotope label in succinate, fumarate, and glutamate after injection with **(A)** ^13^C-glucose or **(B)** U-^13^C-palmitate. Mean ± SEM are shown. Statistics were calculated using one-way ANOVA and post-hoc Dunnett’s test for multiple comparisons to a control group. N=4-5. *p<0.5, **p<.01, ***p<.001, ****p<.0001.

## METHODS

### Animal model

Male C57BL/6J (#000664) mice (7-10 weeks old) from The Jackson Laboratory were used for all animal experiments. Mice were housed in normoxic or hypoxic conditions in the Gladstone Institutes animal facility. Hypoxia was simulated in chambers by mixing N_2_ (Airgas), O_2_ (Airgas, Praxair), and room air using gas regulators. F_i_O_2_ and CO_2_ levels were checked daily and continuously monitored wirelessly. To inhibit the accumulation of CO_2_, soda lime (Fisher Scientific) was added to each chamber. Mouse experiments were approved by the Gladstone Institutes Institutional Animal Care & Use Program (IACUC).

### Open field test

Nitrogen gas was mixed into a 20” x 20” Photobeam Activity System (PAS)-Open Field Chamber (San Diego Instruments) to bring the F_i_O_2_ to the appropriate level. An O_2_ gas sensor (Vernier) was used to monitor the oxygen levels in the chamber. Mice housed at 21%, 11%, and 8% F_i_O_2_ were moved into the chamber of matching F_i_O_2_ for 10 minutes. Their movement was recorded in real time with built-in photobeams.

### Pole test

3-4 days prior to the test, mice were pre-trained on the 50 cm pole. Mice were placed at the top of the pole, facing up and facing down, and the time required to descend from this starting position to the bottom of the pole was measured. Each test was conducted 3 times, and the average score was calculated. For the facing down starting condition, latency to descend was scored as follows: for mice sliding <20% of the pole, the lesser of 1.25 times the total time or 15s was recorded; for mice sliding 20-50% of the pole, 15s was recorded; for mice that fell, a max score of 20s was recorded. For the facing up starting condition, latency to descend was scored as follows: for mice sliding <20% of the pole, 1.25 times the total time was recorded; for mice sliding 20-50% of the pole, 30s was recorded; for mice sliding 50-75%, 45s was recorded; for mice sliding > 75%, 60s was recorded; for mice that fell, a max score of 75s was recorded. All experiments were conducted in a glovebox matching the appropriate F_i_O_2_.

### Blood chemistries

Approximately 90 μL tail blood was collected from mice in a glovebox controlling the F_i_O_2_. Blood was collected into K2EDTA tubes (BD) and analyzed on an iSTAT handheld analyzer (Zoetis) using CHEM8+ cartridges (Zoetis).

### Body temperature measurements

Mouse body temperatures were measured using an infrared camera (FLIR). Mice were imaged and the maximal observed body temperature was recorded. Temperatures were measured in a glovebox controlling the F_i_O_2_.

### Weight and food intake measurements

Mice were weighed daily for the first week of treatment and every 2-4 days after the first week. Food intake was calculated based on the change in weight of food in cages between measurements.

### Hormone measurements

Tail blood was collected from mice in a glovebox controlling the F_i_O_2_ and stored in K2EDTA tubes (BD). Plasma was extracted by centrifuging the blood samples at 2,000 rpm for 10 minutes at 4C and collecting the supernatant. Plasma samples were shipped on dry ice to the UC Davis mouse metabolic phenotyping core. Hormone concentrations were measured using multi-spot assay (insulin, leptin; Meso Scale Discovery), radioimmunoassay (glucagon; Millipore) (corticosterone; MP Biomedicals), or ELISA (ghrelin; Millipore).

### Blood glucose measurements

Blood glucose levels were measured using a handheld glucometer (AimStrip Plus). For fasted measurements, mice were fasted overnight for 12 hours.

### Radiotracers

FDG was prepared by the UCSF Radiopharmaceutical Facility. For FTP synthesis, Methyl 16-bromo-4-thia-hexadecanoate precursor (16-[^18^F]fluoro-4-thia-hexadecanoic acid) was synthesized from 1,12 dibromododecane following previously reported methods (DeGrado et al., 2000). FTP was prepared by reacting the methyl hexadecanoate precursor with [^18^F]fluoride ion followed by saponification of the methyl ester and HPLC purification.

### Mouse positron emission tomography and gamma counting

Mouse PET/CT scans (Inveon, Siemens Medical Solutions) were conducted at the Preclinical Imaging Core facility at China Basin in the UCSF Department of Radiology & Biomedical Imaging. Mice were transported in boxes and transportable N_2_ and O_2_ tanks (Airgas) were used to maintain the appropriate F_i_O_2_ in transit. Mice were anesthetized with isoflurane and administered FDG or FTP via tail vein injections. After 30 minutes, the mice were anesthetized and positioned for PET/CT scan. After completion of the scan, mice were euthanized and organs were harvested. Organs were either imaged using the microPET/CT scanner or entered into a gamma counter (HIDEX Automatic Gamma Counter) for quantification of the radioactivity, which was normalized based on organ weight and dose at scan-time. The percent injected dose per gram was calculated for blood and tissues. Dose at the time of scanning or counting was decay-corrected using the known half-life of the radionuclide used for imaging according to the following equation, with Δ*t* representing the time between injection and scan:

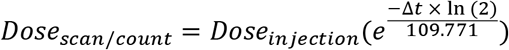

All measured values from microPET/CT and gamma counting were calibrated to quantitatively represent physical activity levels using routine calibration methods employed for both.

### *In vivo* ^13^C-labeled glucose and palmitate tracer experiments

Mice were injected with approximately 1mg/g U-^13^C-glucose (Fisher) or .018 mg/g U-^13^C-palmitic acid (Sigma Aldrich). U-^13^C-glucose doses were dissolved in water and U-^13^C-palmitic acid was conjugated with bovine serum albumin (BSA, 35% water solution) with a PA:BSA molar ratio of 2.65:1. Mice were briefly anesthetized with isoflurane and delivered 100 μL of the tracer through retro-orbital injections. After 20, 45, or 90 minutes, blood was collected using cardiac puncture and stored in K2EDTA tubes (BD). Then, mice were then sacrificed, and organs were harvested and flash frozen in liquid nitrogen.

### Metabolite extraction

Plasma was extracted from blood by centrifuging at 2,000 rpm for 10 minutes at 4C and collecting supernatant. Metabolites were extracted by adding 65 μL of 80% methanol to every 5 μL of plasma. Samples were vortexed for 10s and incubated at 4°C for 10 minutes. Next, samples were centrifuged at 16,000g for 10 minutes at 4°C and supernatant was collected.

For organs, 10 μL methanol and 8 μL water were added for every 1 mg of tissue harvested. A stainless-steel bead (Qiagen) was added to each tube, and samples were lysed by undergoing five 30s on-off cycles in a tissue lyser (Qiagen) at a frequency of 30 Hz. Next, beads were removed and 10 μL chloroform was added for every 1 mg of tissue harvested. Samples were centrifuged at 14,000 rpm for 10 minutes, and the top layer (aqueous) was stored.

### LC-MS

Samples were run on an Orbitrap Exploris 240 high resolution mass spectrometer (HRMS, Thermo Fisher Scientific) using electrospray ionization. The OE240 was coupled to hydrophilic interaction chromatography on a Vanquish Horizon ultra-high performance liquid chromatography (UHPLC, Thermo Fisher Scientific). 5 μLof polar metabolite samples were injected into the LC-MS system and separated on a BEH-Amide column (Waters, 2.1mm x 150mm). The autosampler was maintained at 4°C and the column was maintained at 45°C during runs. Mobile phase A was 20 mM ammonium acetate in 95:5 water:acetonitrile, with ammonium hydroxide added to reach a pH of 9.45. Mobile phase B was acetonitrile. Flow rate was 150 μL/min. The gradient was the following: 0-20 min: 80% B to 20% B; 20-20.5 min: linear gradient from 20-80% B; 20.5-28 min: stable at 80% B (Spinelli et al., 2021). A 3-minute equilibration phase was included at starting conditions before each injection. The OE240 ran in full-scan, negative mode with an ion voltage of 3.0 kV, and the scan range was 70-1000 m/z. The orbitrap resolution was 60000, RF Lens was 50%, AGC target was 1e6, and the maximum injection time was 20 ms. The sheath gas was set to 40 units, auxiliary gas was 15 units, and sweep gas was 1 unit. The ion transfer tube temperature was 275°C, and the vaporizer temperature was 320°C.

## QUANTIFICATION AND STATISTICAL ANALYSIS

### LC-MS data analysis

Peaks were detected and peak areas were quantified using TraceFinder 5.1 General Quan. Natural isotope abundance correction was conducted using the R package AccuCor (Su et al., 2017). Next, relative enrichment of the isotope label was calculated for each metabolite by determining the proportion of carbons that were ^13^-C-labeled. No isotopomers of TCA metabolites were excluded from analysis because all can plausibly emerge from oxidative metabolism of circulating fuels (Fig S3A). For all organ data, relative enrichments of TCA metabolites were normalized to the relative enrichment of circulating glucose or palmitate (Fig S4B). The resulting normalized relative enrichment values were plotted and analyzed using GraphPad Prism. Statistical significance was determined using one-way ANOVA and post-hoc Dunnett’s test for multiple comparisons to a control group.

### Statistical analysis

For all other data analysis and representation, R was used. Graphs were made using ggplot2 (Wickham, 2016). Statistics were calculated using two-way ANOVA and post-hoc Tukey test for multiple comparisons. Statistical significance was depicted on graphs using ggsignif (Ahlmann-Eltze and Patil, 2021). Where used, boxplots depict the median, first quartile, and third quartile.

### Figures

Figures were assembled in Adobe Illustrator. Diagrams were created using Illustrator or Biorender.com.

